# Task context modulates feature-selective responses in area V4

**DOI:** 10.1101/594150

**Authors:** Dina V. Popovkina, Anitha Pasupathy

## Abstract

Feature selectivity of neuronal responses in primate visual cortex is typically measured while animals fixate a small dot on the screen and a stimulus is presented in the near-periphery. This paradigm allows the efficient exploration of feature space, but it provides only a partial view of selectivity by failing to characterize how cognitive factors influence neuronal tuning. Here we focus on primate area V4, known to be influenced by cognitive processes, and ask how neuronal tuning is modulated by task engagement. We compared the tuning for shape and color in 83 well-isolated V4 neurons measured during passive fixation and during active engagement in a shape discrimination task. In both tasks, animals saw the same set of objects—shape x color combinations—but while neither stimulus feature was relevant during the fixation task, shape identity was relevant for behavior during the discrimination task. Consistent with attentional studies, V4 responses during the discrimination task showed a stimulus-independent gain scaling relative to passive fixation, but this was only in a minority of neurons (21/83). For the rest (62/83), response modulations during discrimination depended on stimulus identity: on stimulus shape in neurons more strongly tuned to shape, and on color in neurons more strongly tuned to color. Overall, this resulted in broader tuning for stimulus color, but not shape, during active task engagement. These results suggest that task context can influence the shape and color selectivity of V4 neurons, and in some neurons this effect is consistent with a change in the width of feature tuning.

## INTRODUCTION

Responses of neurons in primate visual area V4 reflect different aspects of visual objects, including form, color, brightness, and texture. Tuning for these object properties has typically been investigated in studies where the animal fixates a central location on the screen, and stimuli are presented in the receptive field (RF) of the neuron. Such characterizations have formed the basis of much of our knowledge about the encoding of visual stimuli throughout the ventral pathway of visual processing, which comprises brain areas involved in object recognition. However, as information about stimuli progresses along the pathway from the primary visual cortex to inferotemporal cortices, neuronal responses display an increasingly prominent influence of cognitive factors (Buffalo *et al*., 2010), which is thought to enable the animal to perform visually-guided tasks.

Many previous studies investigated the influence of cognitive factors in area V4 in the context of attentional paradigms (*e.g.* Haenny *et al*. 1988; McAdams and Maunsell 1999, 2000). These experiments generally compare responses to stimuli within or outside the neuronal RF as the animal was cued to attend to a spatial location and/or an object feature (*e.g.* detect the blue object or the dots moving upward). In these contexts, V4 neurons tend to display enhanced responses to the attended stimulus location or feature. In studies where response magnitude was examined alongside feature selectivity, tuning during the “attended” condition could be explained by applying a single multiplicative factor to the responses during the “unattended” condition (*e.g.* McAdams and Maunsell, 1999). Studies of V4 responses using visual search tasks have reported a similar effect of behavioral goals on responses of V4 neurons, which are influenced by the target feature (*e.g.* Ogawa and Komatsu, 2004). However, in these paradigms the animals perform the same behavioral task while attention is directed to different features of the visual stimulus (e.g. spatial location, contrast, orientation). It is largely unknown whether object representation, as we understand it from fixation studies, is similar when animals are engaged in an active behavioral task - how does selectivity for object features, *e.g.* form and color, compare in these two contexts?

Previous studies of top-down effects on V4 responses suggest that responses during behavior could simply be scaled versions of responses during fixation, and stimulus preference is maintained. However, task context could induce a change in feature selectivity, including a shift in the tuning peak (indicating a change in the neuron’s preferred feature) or changes in the tuning width (indicating an increase in discriminability between preferred and non-preferred features if tuning narrows, or the converse if it broadens). Such changes have been observed in area V4 for comparisons of easy *vs*. difficult discrimination tasks (Spitzer *et al*., 1988) and during feature-based attention (David *et al*., 2008). Feature-based changes have also been reported in the dorsal stream of visual processing, where area LIP (lateral intraparietal area) neurons display shifts in selectivity for stimulus color and direction of motion, depending on the feature relevant for the task (Ibos and Freedman, 2014).

To understand how task context affects stimulus preferences in area V4, here we examined shape- and color-selective responses of 83 V4 neurons to the same stimuli shown in either a passive (fixation) or an active (discrimination) task context.

## METHODS

### Surgical methods

We implanted 2 adult male macaque monkeys (*Macaca mulatta*) with custom headposts and chambers positioned over dorsal area V4 (left hemisphere). The placement of the chamber over the prelunate gyrus and subsequent craniotomy were guided by structural MRI (for details, see Bushnell *et al.,* 2011a). All animal procedures conformed to the National Institutes of Health guidelines and were approved by the Institutional Animal Care and Use Committee at the University of Washington.

### Visual stimulus presentation

Stimuli were presented against a uniform gray background (luminance 5.4 cd/m^2^) using a spectrally calibrated (PR650, PhotoResearch) CRT monitor positioned 45.4 or 56 cm away (subjects 1 and 2, respectively). The animal fixated a small white dot in the center of the screen within a window of radius 0.75°-1°. Eye position was monitored using an infrared eye tracking system (EyeLink 1000; SR Research) and coordinated with stimulus presentation using custom software based on Pype (originally developed by Jack Gallant and James Mazer; Mazer, 2013).

### Task design

#### Fixation task

Each trial of the fixation task (see schematic in **Fig. 1A**) began with the presentation of a fixation spot. Once the animal acquired fixation, a sequence of 3-5 stimuli were presented with a 200 ms blank interval preceding each stimulus. For the first 12 neurons we recorded, each stimulus was presented for 300 ms; for the rest, this presentation time was shortened to 250 ms to better match the timing in the discrimination task (see below). A total of nine stimuli (3 shapes x 3 colors) customized to the shape and color preferences of the neuron under study (see Stimulus Selection, below) were used to probe responses during the fixation task. Stimuli were presented in random order, and each was shown 20 times within a block of fixation task trials. The animal was rewarded with drops of juice for successful fixation for the entire duration of a trial (typically 2000-2500ms).

**Figure 1.**
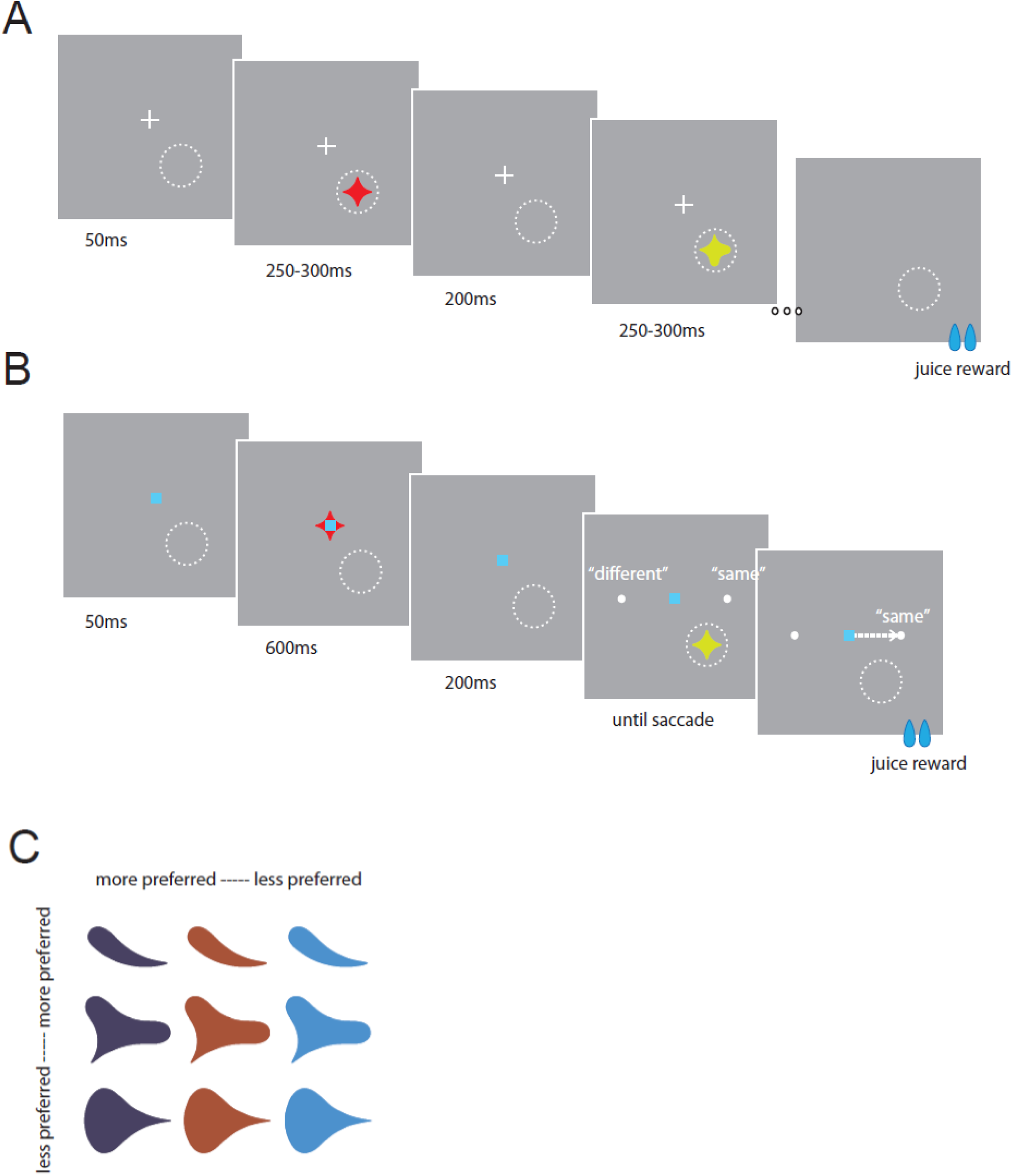
Task conditions and stimulus design. (A). Fixation task. The white dotted circle represents the spatial extent of the receptive field (RF) of the V4 neuron under study and was not presented on the screen. The animal fixated a small white cross at the center of the screen while stimuli were presented in the V4 neuron’s RF one at a time and separated by 200 ms. The animal was rewarded with drops of juice for continuously fixating a sequence containing 3-5 stimuli. (B). Sequential shape discrimination task. The white dotted circle represents the spatial extent of the V4 neuron under study and was not presented on the screen. The animal fixated a small blue square at the center of the screen. Then, a reference stimulus was presented at the center of the screen for 600 ms, followed by a 200 ms interval. A test stimulus was then presented in the V4 neuron’s RF along with two saccade choice target dots; the animal then had 1.5s to report whether the shape of the test stimulus matched the reference stimulus with a rightward (match) or leftward (non-match) saccade. The animal was rewarded with drops of juice for a correct saccade. In the example trial shown here, the two stimuli had the same shape so the animal would be rewarded for a saccade to the rightward target, represented by arrow. On average, animals took 260 ms to report their choice. (C). Example stimuli from one session. We chose 3 shapes that spanned the dynamic range for each neuron based on preliminary screening; i.e., we chose a shape that elicited a strong response, a weak response, and an intermediate level response. We used the same process to choose 3 colors (differing in hue and luminance), and created a set of 9 stimuli with unique shape-color pairings that were then presented in the task contexts schematized in (A) and (B). Colors shown are adjusted for visibility in print.

#### Discrimination task

Animals were also trained on a sequential shape discrimination task (**Fig. 1B**). Each trial began with the presentation of a fixation spot. Once the animal acquired fixation, a “reference” stimulus was presented at central fixation for a duration of 600 ms. After an inter-stimulus interval of 200 ms, a “test” stimulus appeared within the receptive field (RF) of the V4 neuron. Simultaneously, two small target dots appeared to the left and right of the fixation spot, each 6 degrees of visual angle away from fixation. The animal then had up to 1500 ms to report whether the shape of the test stimulus was the same or different from the reference stimulus via a saccadic eye-movement to the right or left target dots, respectively. The test stimulus disappeared as soon as the animal’s eye left the fixation window; test stimulus duration varied from trial to trial, dictated by the reaction time of the animal. Mean reaction times, measured as the time at which the eye exited the fixation window, were 260±23 ms across both animals (M1: 239±21 ms; M2: 272±12 ms). The same 9 stimuli used during the fixation task served as the reference and test stimuli in the discrimination task. Each of the 9 stimuli appeared randomly as a test stimulus for a total of 15-30 repeated presentations in each block of discrimination trials. The animal was rewarded with drops of juice for a saccade to the correct target location; the number of trials requiring a leftward or rightward saccade were balanced and chance performance was 50%; mean animal performance was 83±5% (M1: 85±7%; M2: 81±3%). In our analyses below, we include only responses to stimuli where the outcome of the trial was a correctly directed saccade.

### Data collection

Each day, we lowered a single tungsten microelectrode (FHC) using a stepper motor drive (Gray Matter Research). We amplified and filtered the signals from the electrode and sorted waveforms (Plexon Systems) to identify single units. Here, we include responses from 83 well-isolated neurons for which RF eccentricities ranged from 1.6° to 7.3°.

#### Stimulus selection

For each neuron in our dataset, we first assessed the spatial extent of the RF manually using a variety of shapes and colors under mouse control. We then characterized each neuron’s color/luminance preference either manually (6 neurons) or via an automated protocol that measured neuronal responses to a single shape presented in 25 colors that uniformly sampled the CIE space, and at 3-4 luminance levels (see Bushnell *et al*., 2011b). To assess shape selectivity, we presented 25 2D shapes (a subset of the shapes used by Pasupathy and Connor, 2001) at 8 rotations, for a total of 200 shape stimuli.

Based on the color/luminance and shape preferences of each neuron, we chose three shapes and three colors that represented the preferred, intermediate and non-preferred values along each of the two feature dimensions for each neuron. We then created a set of 9 stimuli by combining the shapes and colors thus selected for use during the fixation and behavioral tasks (see Figure 1C for an example set from one session). When responses of more than one neuron were recorded simultaneously (14 sessions in animal 2), stimuli were tailored according to the preferences of only one of the neurons. Given evidence that shape selectivity in V4 is independent of stimulus color (Bushnell et al. 2012), our choice of stimuli enabled us to consider the influence of task on tuning separately for shape and for color.

Position of the test stimuli during behavior and all stimuli during fixation were jittered by up to 5 pixels about the center of the neuron’s RF. Stimuli were scaled such that all parts of the stimuli were within 80% of the estimated RF diameter based on data in Gattass *et al*. (1988); reference stimuli in the discrimination task were size-matched to the RF-based scaling of the test stimuli.

For each neuron, fixation and behavioral tasks were conducted in alternating blocks starting with the fixation task. Each fixation block was 45-60 trials long, and each discrimination block was 144-288 trials long; as described above, 3-5 stimuli were presented per fixation trial, while one stimulus was presented in the RF per behavior trial. The characteristics of the fixation spot identified the task: a white cross indicated a fixation task while a blue square indicated a discrimination task. Here we include responses from neurons where we successfully captured at least 2 blocks of both tasks (median: 3 blocks for each task).

### Data Analysis

#### Computing neuronal responses

Stimulus onset and offset times were detected with a photodiode. To construct peristimulus time histograms for each neuron (PSTH, as in **Fig. 2A**), responses in all blocks of one task type (fixation or discrimination) were aligned to stimulus onset and 1-ms bin responses were smoothed with a Gaussian kernel (σ = 10 ms). To compute average responses for each stimulus, we computed the mean spiking rate on each trial in the time window of 50-200 ms from stimulus onset. This time window was chosen to account for V4 response latency (Schmolesky *et al*., 1998; Zamarashkina *et al*., 2017). The later bound of 200 ms was chosen to discount the influence of reward anticipation or saccade preparation (in the discrimination task, time to saccade initiation was 260±23 ms). To construct stimulus-response (S-R) curves (as in **Fig. 2B**), we further averaged responses across multiple presentations of each stimulus.

**Figure 2.**
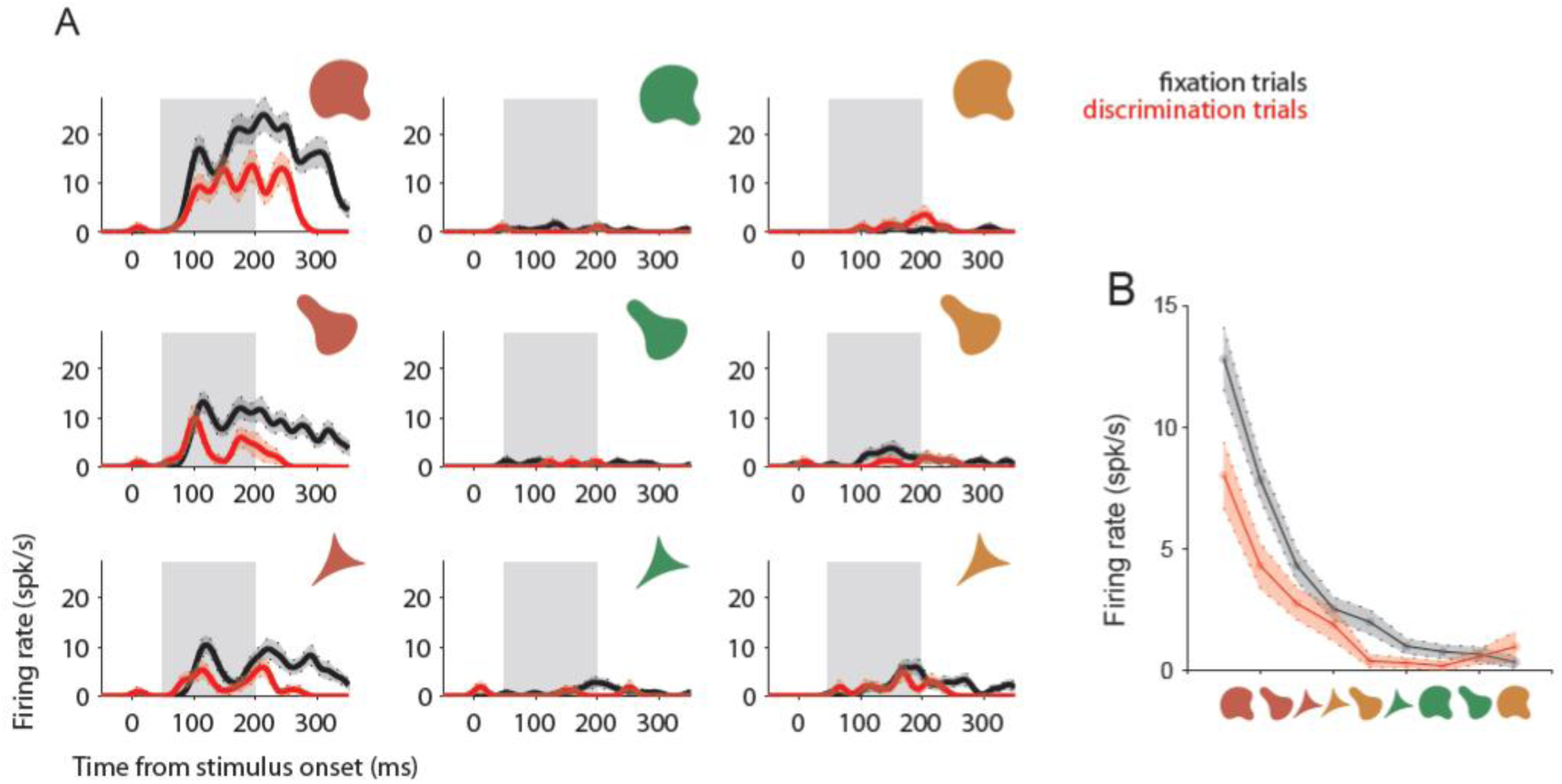
Responses of example neuron 1 (p150514) to stimuli in fixation and discrimination task conditions. (A). Peristimulus time histograms (PSTH) for responses of neuron #1 to 9 stimuli while the animal performed a fixation task (black traces) or discrimination task (red traces). The corresponding stimulus for each panel is shown in the insets; panels are arranged such that stimuli sharing the same shape are in rows, and sharing the same color are in columns. Responses of neuron #1 were both color- and shape-selective during the fixation task (compare responses to stimuli in top row; stimuli in left column). During the discrimination task, responses remained shape- and color-selective, and response magnitude decreased for all stimuli. Gray boxes indicates time window for response averaging (see below). (B). Schematic representation of tuning for neuron #1. Each point represents the average response to the stimulus depicted along the abscissa (50-200 ms time window, see gray window in A). Abscissa is ordered according to stimulus preference, *i.e.* response magnitude during the fixation task (black trace), and responses during the discrimination task shown in red. Upper and lower bounds are SEM. Note the similar shape of the tuning schematic for the two task conditions.

To obtain feature-specific average PSTHs (*i.e.* shape- or color-specific responses) we first averaged across all stimuli with a particular feature, *e.g.* the preferred shape in all colors. Each feature-specific PSTH thus contained 3 responses: one for the preferred feature, one for the intermediate, and one for the nonpreferred. We normalized these PSTH traces separately for each task and neuron, and averaged the resulting PSTHs across neurons. Finally, we obtained tuning curves, as above, by computing average responses in the 50-200 ms window for each of the 3 stimuli, and normalized them such that the response to the preferred feature was 1.

#### Regression models

To compare neuronal responses to the 9 stimuli during fixation and discrimination, we used individual trial responses to fit four regression models. Based on the body of literature examining attentional influences in V4, we expected to see a difference in responses during the discrimination task, and chose a variety of models to compare our observations to previously published results.

First, we considered the possibility that neuronal responses during the behavioral task scaled linearly relative to the responses during the fixation task, as reported in attentional studies; such a response change could be described using the linear-gain model:

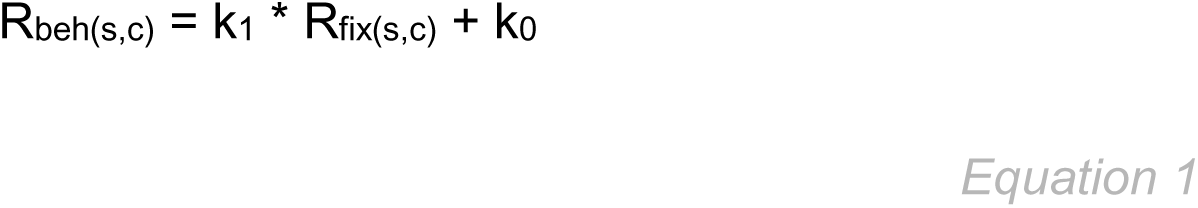

where R_beh_ and R_fix_ denote the neuronal response during the discrimination and fixation tasks respectively, and k_1_ and k_0_ are the slope and constant for the regression.

To perform the regression, we needed to pair trials from fixation and discrimination blocks (R_fix_ and R_beh_). Fewer trials were shown in the discrimination condition; thus, we first resampled responses during the discrimination task using a bootstrap procedure, randomly drawing responses (with replacement) to match the number of trials during the fixation task. We then randomly paired responses during the fixation task with responses during the discrimination task for each of the 9 stimuli (R_fix(s,c)_ and R_beh(s,c)_). To reduce the potential introduction of bias, we repeated the resampling and pairing procedure to obtain 100 sets of regression data, and performed subsequent model fitting and analyses of model performance across these sets.

Additionally, we considered a model where neuronal responses during the discrimination task also depend on the responses during the fixation task, but this relationship is non-linear (polynomial-gain model):

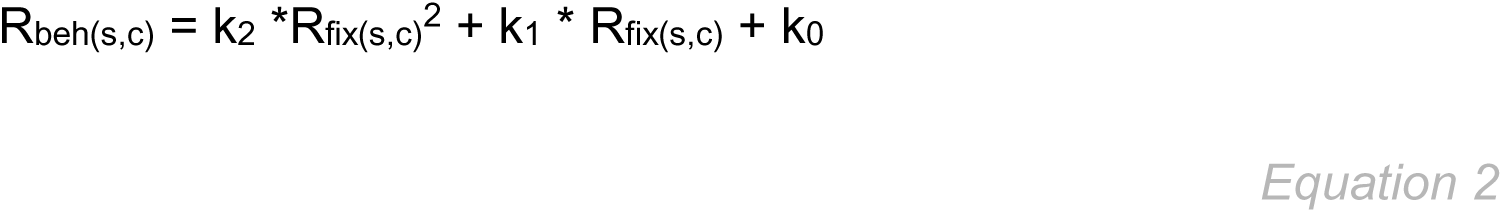

where k_2_ and k_1_ are the 2^nd^ and 1^st^ degree regression coefficients, and k_0_ is the regression constant. We also considered exponential and logarithmic forms of this nonlinear relationship, but did not find these models to fit our data better than the 2^nd^ degree polynomial form.

As an alternative to firing-rate based models, we considered a case where neuronal responses during the discrimination task depend not only on responses during the fixation task, but also on the features of the stimulus in its RF. We defined feature-gain models for shape and color of the stimulus, as follows:

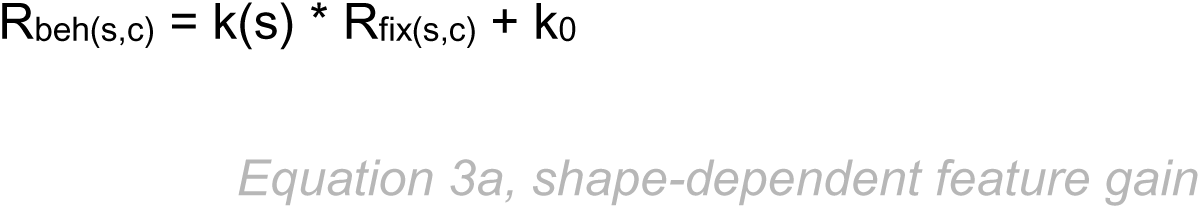

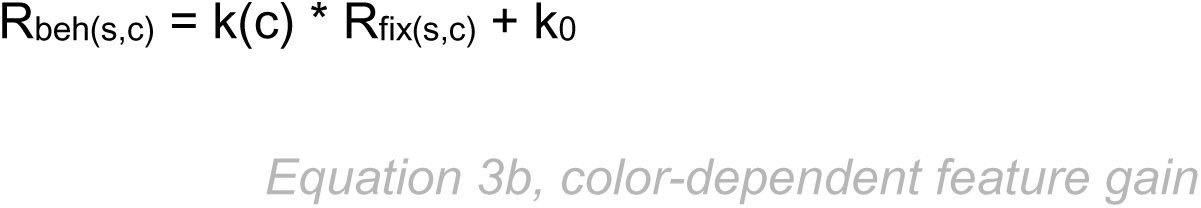

Where k(s) and k(c) represent separate coefficients for each of 3 stimulus shapes or colors, respectively, and k_0_ is the regression constant.

#### Model comparison

To compare the models above, which have different numbers of parameter constraints, we performed cross-validation using 4/5 of the neuronal data to fit each regression model, and the remaining 1/5 to test the model fit. Responses to each of the 9 stimuli were included in both the model fitting and testing sets. As described above in *Regression models*, we repeatedly randomized the pairing of responses during fixation and discrimination tasks; we performed this cross-validation procedure for each random pairing.

To quantify performance, we calculated root mean squared error (RMSE) for the test sets, and compared the RMSE distributions and average RMSE values for each model to determine the best-fitting model for each neuron. A model was determined to fit better if its average RMSE value was lowest and the distribution of RMSE values was significantly different than other models (unpaired t-test, α=0.01; computed for pairs of models).

To obtain predicted responses during discrimination using each regression model, we first averaged model coefficients over 100 randomized pairings of fixation-discrimination trials. We then used the averaged coefficients together with the average responses to 9 unique stimuli in the fixation condition to generate predicted discrimination responses from Eqns. 1-4.

#### Feature selectivity

For each neuron, we estimated the strength of feature selectivity for shape and color by calculating the partitioned variance due to each feature from a 2-way ANOVA model with shape, color, and interaction terms. The selectivity metric, η^2^, was calculated as the partitioned sum of squares for each factor (shape, for η^2^_shape_; or color, for η^2^_color_) divided by the total sum of squares. The interaction term was small for all neurons; on average, it accounted for 4% of the relative variance.

To compare the shape or color bias in feature selectivity of individual neurons, we computed a ratio of η^2^_shape_ to η^2^_color_. This metric is >1 for more shape- than color-selective neurons, and <1 for more color- than shape-selective neurons.

#### Contribution of other task-dependent variables

To determine whether the identity of the reference stimulus contributed to the observed effects, we first separated neuronal responses during the discrimination task into two sets according to whether the reference stimulus matched or did not match the neuronal preference for shape and color. The first set of data contained responses to test stimuli in trials where the reference stimulus was of the preferred shape and color (reference-preferred), and the second set contained responses to test stimuli in trials where the reference stimulus had any other combination of shape and color (reference-other).

We then examined the magnitude of the reference stimulus-dependent effect, captured in the difference between responses in the reference-preferred and reference-other datasets, relative to the overall responsiveness of the neuron using the following ratio:

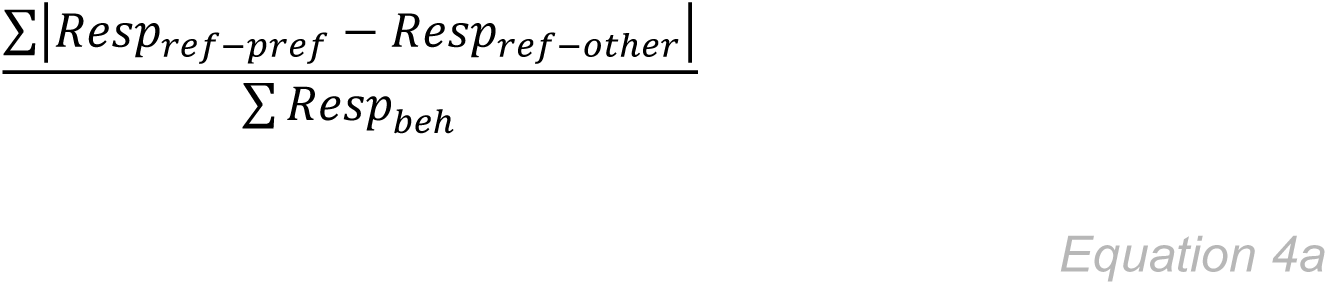

where Resp_ref-pref_ is the mean response for each of 9 test stimuli for the subset of data where the reference stimulus had the preferred shape and color, Resp_ref-other_ is the mean response to each of the 9 test stimuli for the subset of data where the reference stimulus had other shape and color combinations, and Resp_beh_ is the average response to each of the 9 stimuli during the discrimination trials. For a neuron with a mean response of 50 spk/s, a ratio with a value of ~1 represents a mean stimulus response difference of 5 spk/s between the two data subsets.

We performed the same analysis to determine whether the motor plan being prepared by the animal in the behavioral condition (a saccadic eye movement) contributed to the effects we observed. We separated neuronal responses during the discrimination task into two sets according to the saccade outcome of the task, and compared the magnitude of the saccade-related effect using an equivalent ratio to Eqn. 4a:

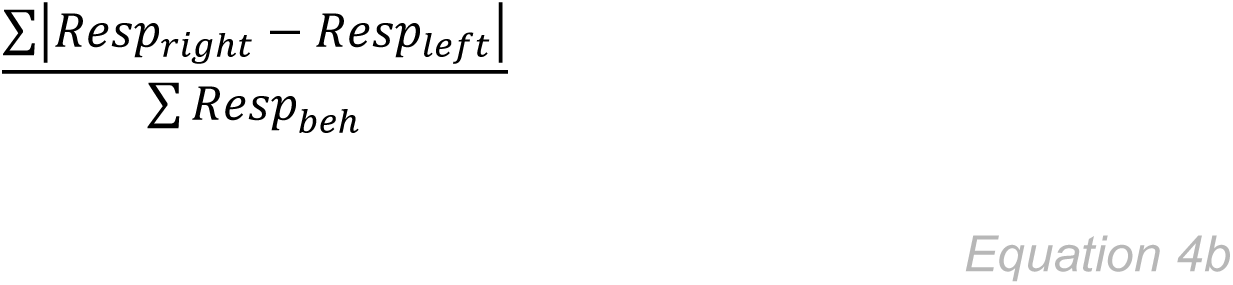

where Resp_right_ is the mean response for each of 9 test stimuli for the subset of data where the trial ended in a rightward saccade, Resp_left_ is the mean response to each of the 9 test stimuli for the subset of data where the trial ended in a leftward saccade, and Resp_beh_ is the average response to each of the 9 stimuli during the discrimination trials. For a neuron with a mean response of 50 spk/s, a ratio with a value of ~1 represents a mean stimulus response difference of 5 spk/s between the two data subsets.

## RESULTS

To examine whether feature tuning in area V4 changes depending on the task being performed, we studied the responses of 83 well-isolated V4 neurons (31 from M1, 52 from M2) to colored 2-D shape stimuli presented in two contexts: a passive fixation task, or an active shape discrimination task. During the fixation task (**Fig. 1A**), a sequence of stimuli was presented within the RF of the V4 neuron while the animal maintained fixation at a central location on the screen. During the discrimination task (**Fig. 1B**), the animal had to judge whether two sequential stimuli had the same or different shape and report the decision with a rightward or leftward saccadic eye movement, respectively.

For each neuron, we selected 9 stimuli which elicited responses that spanned the dynamic range of the neuron’s firing rate. Stimuli were composed from combinations of three 2-D shapes and three different colors (see Methods for details; sample stimuli from one session are shown in Figure 1C).

### Task-driven changes in stimulus preference

First, we examined whether task identity modulated stimulus preference in V4 neuron responses. Below, we include three representative examples to demonstrate how V4 response magnitude and feature tuning changed during the discrimination task, compared to the fixation task.

The example neuron in Figure 2 (Neuron 1) displays suppressed responses during the discrimination task, but no change in shape and color preferences. During the fixation task (black traces), responses of this neuron were both shape- and color-selective, i.e. modulated both by stimulus shape (*cf.* PSTHs for stimuli in top row) and color (*cf*. PSTHs for stimuli in left column). The stimulus in the top left panel elicited the strongest responses from this neuron during fixation. During the shape discrimination task, firing rate was suppressed relative to responses during fixation (*cf.* red PSTH traces to black PSTH traces). Neuronal responses retained shape and color selectivity, and the stimulus in the top left panel continued to elicit the strongest responses.

To summarize changes in response magnitude and stimulus preference, we created a stimulus-response (S-R) curve. Unlike stimuli varying in a single feature dimension such as orientation or contrast, our shape-color combination stimuli do not lie on a naturally ordered axis. To visualize the S-R curve for neurons under study, we created a custom axis by ordering stimuli based on the magnitude of responses during the fixation task, which we averaged in the 50-200 ms window after stimulus onset (gray boxes in **Fig. 2A**). Figure 2B shows the S-R curve for Neuron 1, with stimuli depicted on the abscissa; black traces represent average neuronal responses during the fixation task, and red traces during the discrimination task. The resulting curve shows that the responses to stimuli during the discrimination task were lower than during the fixation task, but tuning was similar. Such a change in responses may reflect simple scaling of responses as shown in studies of attention (*e.g*. McAdams and Maunsell, 1999).

Unlike what we observed for neuron 1, in other neurons we observed changes in responses during the discrimination task that did not preserve ranked stimulus preference. Figure 3A shows the responses of one such neuron. Like responses of Neuron 1 above, when the animal performed the fixation task (black traces), this neuron’s responses were modulated both by stimulus shape (*cf*. PSTHs for stimuli in top row) and color (*cf*. PSTHs for stimuli in left column). The stimulus in the top left panel elicited the strongest responses during fixation. During the discrimination task, responses of Neuron 2 increased (red traces), but the changes in firing rate were not uniform across stimuli. That is, responses to some stimuli (e.g. top right) increased more than others. Two points are notable: the most preferred stimulus (top left) was the same in the two tasks; and stimuli with the same shape (*e.g.* panels in top row) elicited a similar magnitude of response during the discrimination task. This observation suggests that for Neuron 2, stimulus shape modulated response magnitude in discrimination trials more strongly than stimulus color.

**Figure 3.**
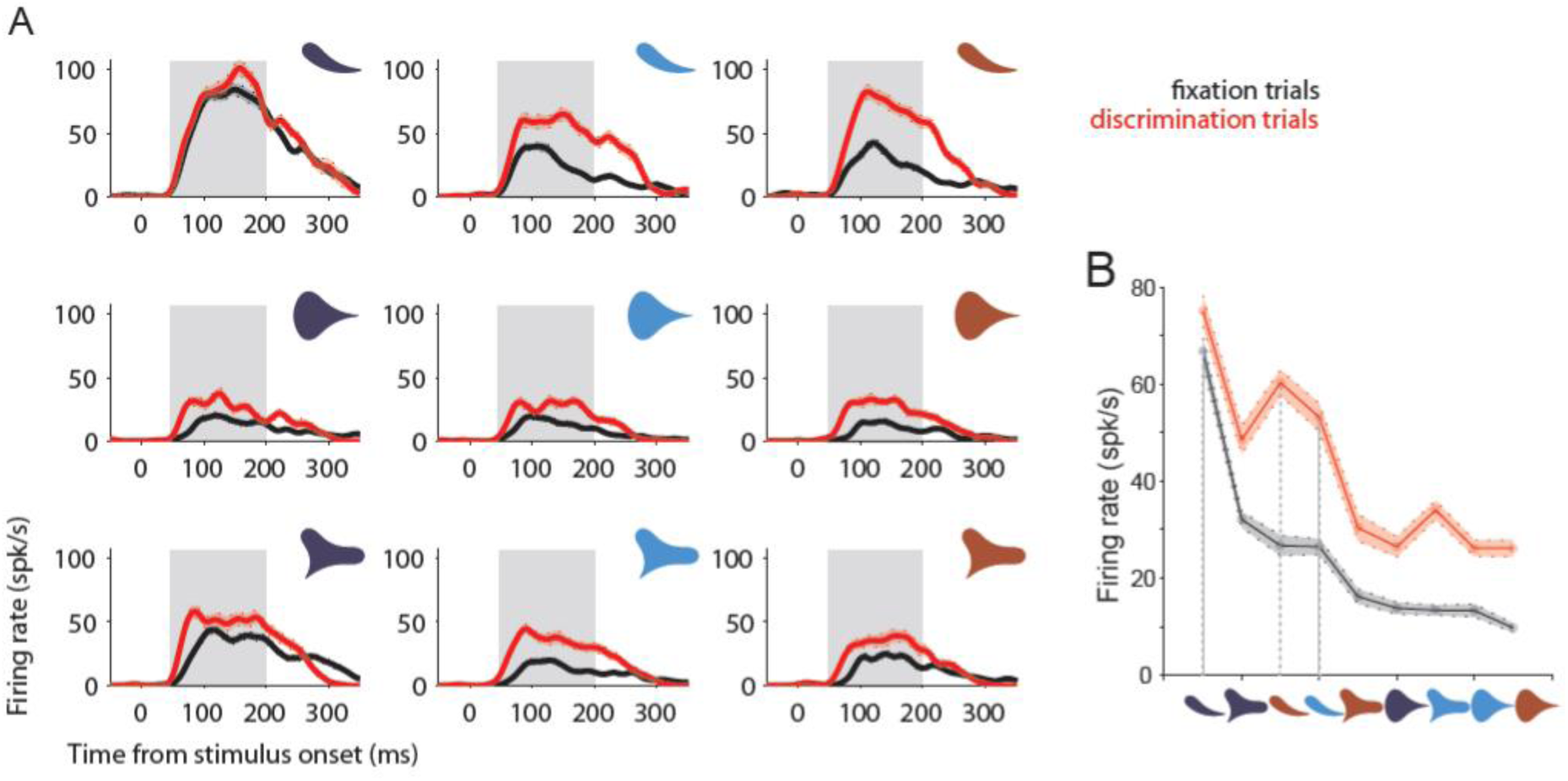
Responses of example neuron 2 (o140605) to stimuli in fixation and discrimination task conditions. (A). PSTH for responses of neuron #2 while the animal performed a fixation task or discrimination task; layout and color scheme as in Fig. 2A. Responses of neuron #2 were both color- and shape-selective during the fixation task (compare responses to stimuli in top row; stimuli in left column, respectively). Note that during the discrimination task, responses to all stimuli with the same shape (in rows) became more similar in magnitude across different colors. Traces are averaged across all blocks of the same task condition; upper and lower bounds represent SEM. (B). Schematic representation of tuning for neuron #2, computed as in Fig. 2B. Grey dashed lines highlight responses to stimuli in top row of (A).

The S-R curve for Neuron 2 (Fig. 3B) illustrates the point that the rank order of responses differed during fixation and discrimination blocks. Specifically, the “tadpole” shaped preferred stimulus (Fig. 3A, top row) in brown and cyan evoked stronger responses during discrimination than would be expected from the responses during fixation (*e.g.*, dashed lines highlight responses of stimuli in top row of **Fig. 3A**). This change in the tuning profile suggests that, unlike in Neuron 1, responses of Neuron 2 during discrimination were not consistent with a simple scaling of responses during fixation.

While data above were averaged across two blocks of the same task, we also examined responses within a single block of trials to ensure the comparisons were robust. In all recording sessions, observed effects were consistent across task blocks. For example, for Neuron 2, the dissimilarity in the ranked stimulus preference during fixation and discrimination tasks persisted across repeated blocks of trials.

Figure 4 presents the responses of an additional neuron where stimulus preference is not maintained. Like the other two examples, this neuron’s responses during fixation trials (black traces, **Fig. 4A**) were modulated by both shape and color (*cf.* PSTHs for stimuli in top row and left column, respectively). The stimulus shown in the top left panel elicited the strongest response from this neuron in both task contexts. While firing rate increased for some stimuli during the discrimination task, it did not for others. This difference is best captured by the S-R curve for Neuron 3 (Fig. 4B). During discrimination trials, red-colored stimuli did not elicit the same increase in FR as the other stimuli, an increase which would have been expected given the ranked stimulus preference during fixation trials (dashed lines highlight responses to stimuli in center column of **Fig. 4A**). This result is complementary to our observations for Neuron 2, suggesting that responses during discrimination were modulated more strongly by stimulus color than stimulus shape.

**Figure 4.**
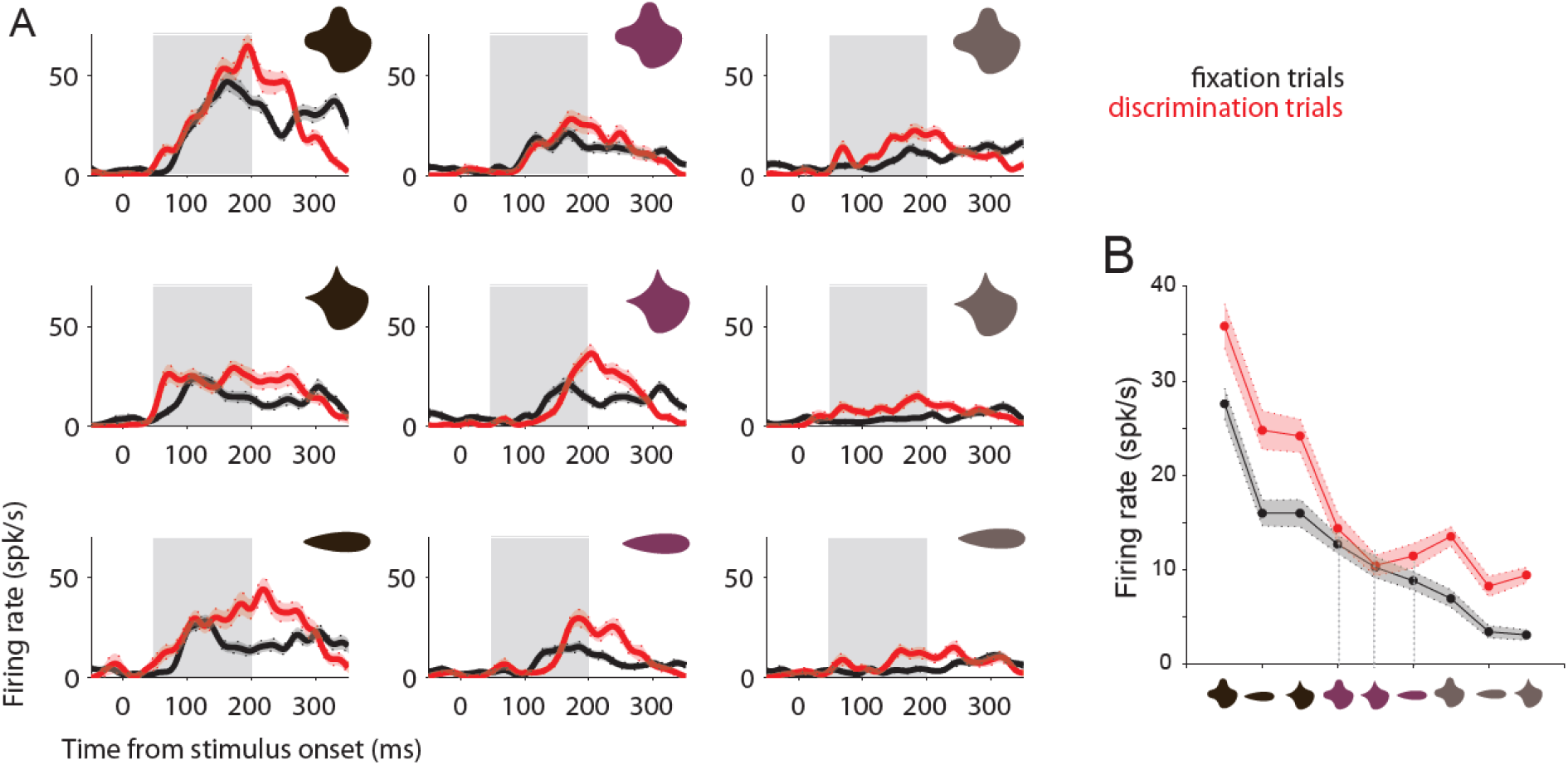
Responses of example neuron 3 (p150409) to stimuli in fixation and discrimination task conditions. (A). PSTH for responses of neuron #3 while the animal performed a fixation task or discrimination task; layout and color scheme as in Fig. 2A. Responses of neuron #3 were both color- and shape-selective during the fixation task (compare responses to stimuli in top row; stimuli in left column, respectively). (B). Schematic representation of tuning for neuron #3, computed as in Fig. 2B. Grey dashed lines indicate responses to stimuli in middle column of (A).

In summary, we observed a variety of changes in responses of individual neurons during fixation and discrimination tasks. In the examples described above, the most preferred stimulus remained the same in both tasks. That is, the response changes in these neurons do not reflect a shift in the peak of tuning for shape or color. We did observe a few neurons where the most preferred stimulus changed, which would be consistent with a peak shift (see Supplementary Figure S1 for an example). However, these few neurons were not representative of the response changes we observed in the population (for further details regarding feature tuning observed in example neurons, see Supplementary Figure S2).

### Modeling firing rate- or stimulus-based scaling to explain modulation during discrimination trials

To further examine the neuronal responses exemplified above, we compared 4 candidate regression models (Eqns 1-3b) to understand what factors could account for the change in responses observed in individual neurons. We considered two models in which predictions were based on scaling the magnitude of responses during fixation trials with fixed factors. These included a linear, single-gain model (Eqn. 1) and a nonlinear, 2^nd^ degree polynomial-gain model (Eqn 2). In the predictions of these models, response magnitude could change during the discrimination trials, but ranked stimulus preference is maintained. This type of modulation would be consistent with observed responses for Neuron 1. We also considered two alternative models in which predictions were based on scaling response magnitude during fixation while taking into account the stimulus features. These included a model with separate gain factors for each stimulus shape (Eqn. 3a) and a model with separate gain factors for each stimulus color (Eqn 3b). In the predictions of these models, both response magnitude and ranked stimulus preference could change during the discrimination trials. This type of modulation would be consistent with observed responses for Neurons 2 and 3.

To fit regression coefficients for each of the models, we used averaged neuronal responses (as for the S-R curves above). For each of the 9 stimuli, we randomly paired fixation and discrimination trials, resampling to equate the number of observations (see Methods for details). Because the models contained a different number of parameters, we used cross-validation to mitigate overfitting. To obtain a robust estimate of RMSE values for model comparison, we repeated the random trial pairing and cross-validation procedure 100 times for each neuron. Below, we compare the average performance of the 4 candidate models: first, for the example neurons, and then, for the entire population of 83 neurons.

**Figure 5** compares the observed (filled) and predicted (open) responses during discrimination trials for example neurons 1-3 (observed: same data as in Figs. 2-4). For Neuron 1, observed responses during discrimination trials were well-captured by the linear-gain model (**Fig. 5A**; mean RMSE = 4.5). The polynomial-gain model and the feature-dependent models (**Figs. 5B-D**) produced similar predictions (mean RMSE: 4.5-4.6), and their performance was not significantly different (unpaired t-test, p > 0.2 for all pairwise model comparisons; α = 0.01). Overall, the linear-gain model provided the most parsimonious explanation of this neuron’s response change during discrimination trials. For Neuron 2, the predictions of the firing-rate dependent models were similar (**Fig. 5 E-F**; mean RMSE: 20.2; unpaired t-test, p = 0.94). The predicted responses of the stimulus color-dependent model (Fig 5H) were similar to the linear-gain model (mean RMSE: 20.2; unpaired t-test, p = 0.73). However, the shape-dependent model captured the responses significantly better (**Fig. 5G**, mean RMSE: 18.4; unpaired t-test, p < 0.001); thus, stimulus shape provided the best explanation for the response modulation displayed by Neuron 2. In contrast, for Neuron 3, the linear-gain, polynomial-gain, and shape-dependent gain models (**Fig. 5I-K**) all produced similar discrimination response predictions (mean RMSE: 11.8-11.9; unpaired t-test, p > 0.03 for all pairwise comparisons). However, the color-dependent model (**Fig. 5L**) explained the observed discrimination responses significantly better (mean RMSE = 10.9; unpaired t-test, p < 0.001 for all pairwise comparisons to the other models); thus, stimulus color provided the best explanation for the response modulation displayed by Neuron 3.

**Figure 5.**
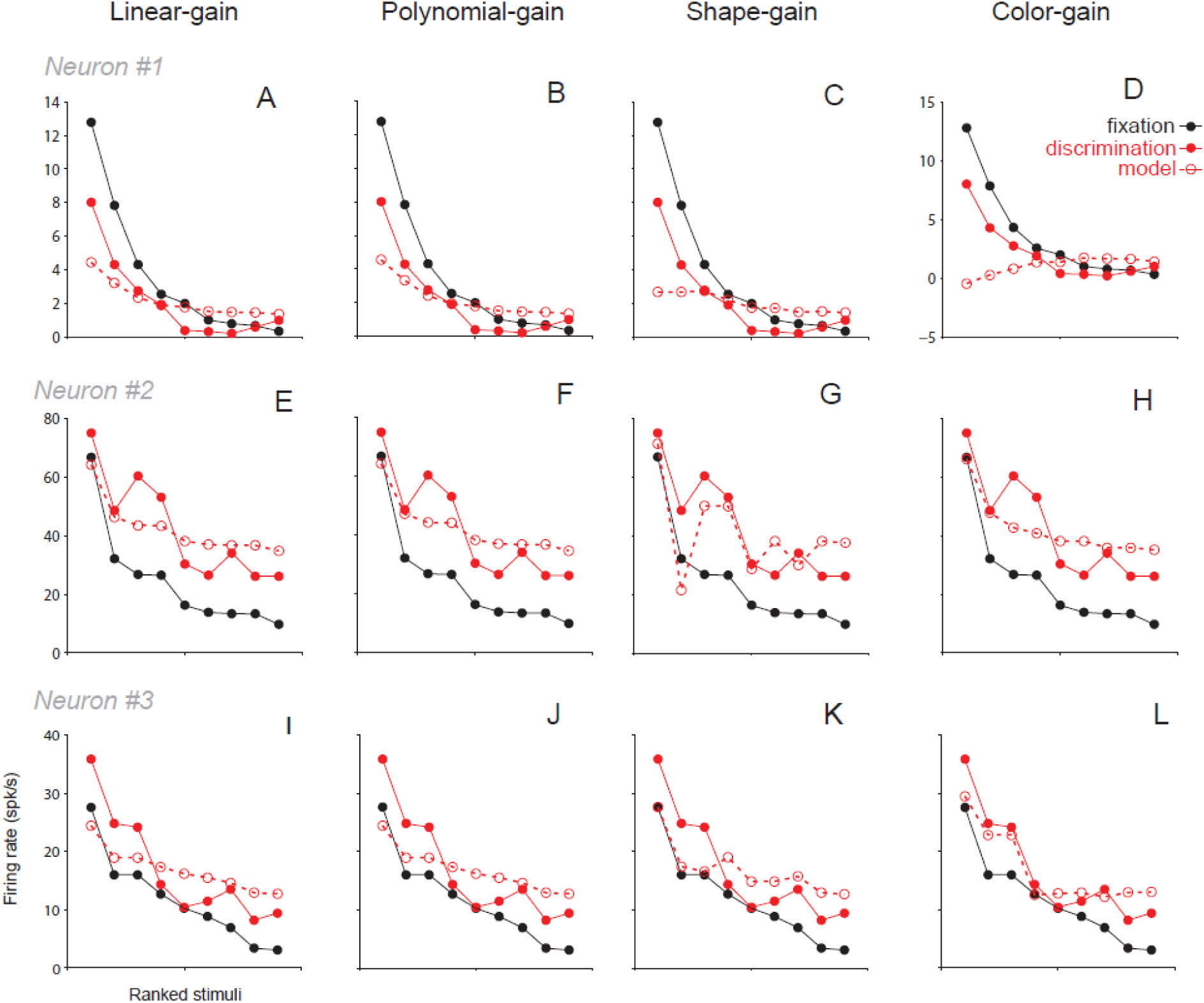
Comparison of predicted model responses for example neurons. (A-D). Observed (red solid line and filled symbols, discrimination task responses) and predicted (red dashed line and open symbols) responses for regression models fit using fixation task responses of neuron #1 (black); discrimination and fixation responses are same data as Fig. 2B. The linear- (A) and polynomial gain models (B) have coefficients that depend on the firing rate of the neuron and produce predictions that match the observed data better than the stimulus-dependent predictions of the shape- and color-dependent gain models (C,D). (E-H). Observed and predicted responses for regression models fit using fixation task responses of neuron #2; discrimination and fixation responses are same data as Fig. 3B. The model in (G) has separate scaling coefficients for stimuli with different shapes, and its prediction matches the observed data best. The rest of the models predict a more uniform scaling of the responses during the fixation task. (I-L). Observed and predicted responses for regression models fit using fixation task responses of neuron #3; discrimination and fixation responses are same data as Fig. 4B. The model in (L) has separate scaling coefficients for stimuli with different colors, and its prediction matches the observed data best. The rest of the models predict a more uniform scaling of the responses during the fixation task.

These examples were representative for our recorded population of 83 neurons. When comparing RMSE values across the 4 candidate models, for 18/83 neurons, the linear-gain model performed best. For 3/83 neurons, the polynomial-gain model performed best. Thus, for 21/83 neurons in our population (25.3%), responses during discrimination could be expressed as a linearly or non-linearly scaled version of the responses during the fixation task. For 20/83 neurons, the shape-dependent model performed best, and for 42/83 neurons the color-dependent model performed best. Thus, for 62/83 neurons in our population (74.7%), response scaling during discrimination was dictated by particular stimulus features.

### Underlying shape or color selectivity correlates with observed modulation

Next, we asked whether the observed response modulation was related to selectivity for stimulus features. Since shape and color preferences are independent in V4 neurons (Bushnell *et al*., 2012), we considered the strength of shape- or color-selectivity across the population, the shape- or color-selectivity bias within each neuron, and tuning width for stimulus shape and color.

#### Strength of feature selectivity

To calculate a proxy metric for the strength of shape or color selectivity, we took each neuron’s responses to the 9 unique stimuli and computed η^2^: the partitioned sum of squares associated with stimulus shape (η^2^_shape_) or stimulus color (η^2^_color_), normalized by the total sum of squares. A higher value for η^2^_shape_ corresponds to more of the response variance captured by shape, and thus stronger shape selectivity; and the converse for η^2^_color_ (also known as ‘effect size’; as a rule of thumb, ~0.13 – medium effect, ~0.26 – large effect). We found a difference in the strength of stimulus feature selectivity that correlated with the best-fitting model for each neuron (Fig 6). The observations were similar regardless of whether the metric was computed using responses during fixation or during discrimination trials. Neurons whose response modulation was best explained by the shape-dependent model (red) were associated with significantly higher η^2^_shape_ than other neurons (unpaired t-test, p << 0.001). Neurons best explained by the color-dependent model and both firing-rate models had similar shape selectivity (p = 0.14). The converse was true for color selectivity. Neurons whose responses were best explained by the color-dependent model (**Fig. 6B**, blue) had significantly higher η^2^_color_ than the other neurons (unpaired t-test, p << 0.001). Neurons best explained by the shape-dependent model and both firing-rate models had similar color selectivity (p = 0.32).

**Figure 6.**
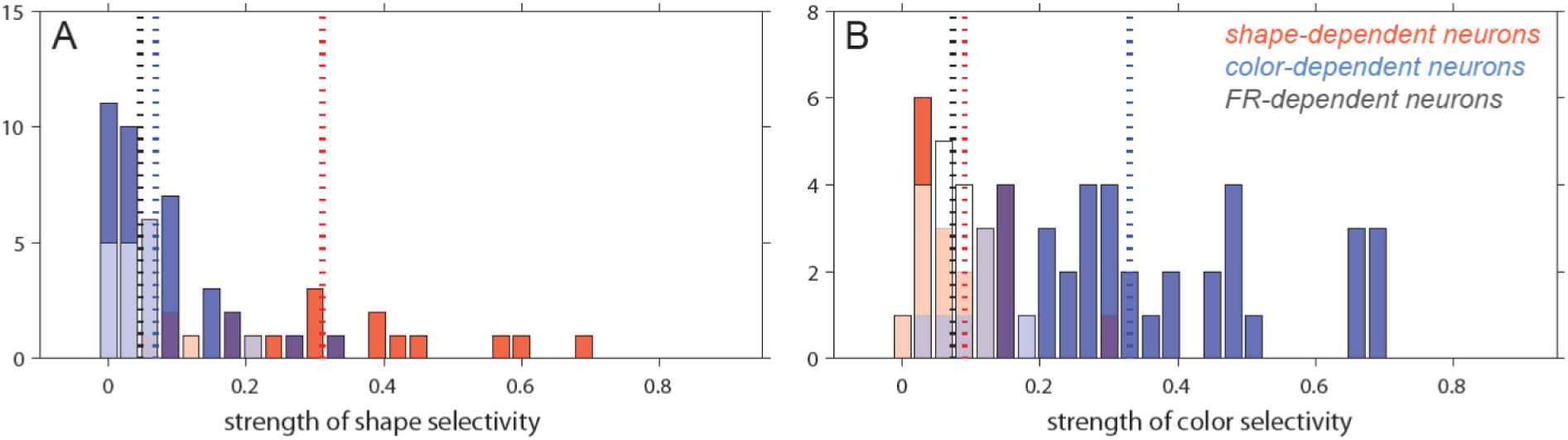
Relationship between individual feature selectivity and best-fitting model. Individual feature selectivity was calculated as the partitioned variance due to shape or color from 2-way ANOVA (see Methods). (A). Comparison between linear-gain and shape-dependent feature gain model. The histograms show the relative strength of shape selectivity for neurons whose responses are better fit by the shape-dependent feature gain model (red) compared to those that are better fit by the linear-gain model (white). Shape and color selectivity computed as partitioned variance from 2-way ANOVA (see Methods). (B). Histograms as in (A), shown for relative strength of color selectivity in neurons whose responses are better fit by the color-dependent feature gain model (blue) compared to those that are better fit by the linear-gain model (white).

#### Feature selectivity bias

Each neuron in V4 could be relatively more shape- than color-selective, more color- than shape-selective, or selectivity could be similar for both features. To understand how bias for shape- or color-selectivity relates to the observed response modulation, we computed a ratio between η^2^_shape_ and η^2^_color_, and compared this value computed for responses during the discrimination task (**Fig. 7**, ordinate) to that computed for responses during the fixation task (abscissa).

**Figure 7.**
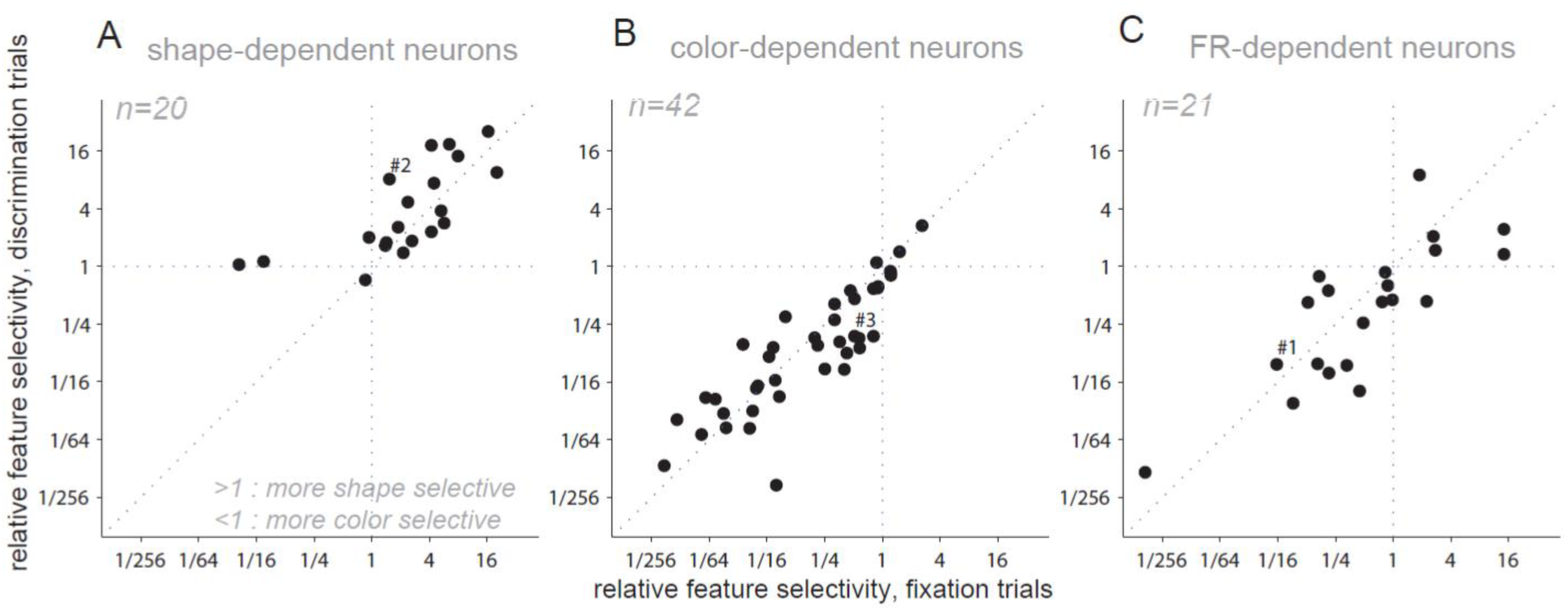
Relationship between selectivity bias and best-fitting model. Selectivity bias, calculated as the ratio between partitioned variance due to shape and color from 2-way ANOVA (see Methods), is shown for discrimination trials (ordinate) compared to fixation trials (abscissa). Examples neurons #1-3 are marked. (A). Selectivity bias for neurons whose responses were well-fit by the shape-dependent model. Relative selectivity of > 1 indicates most responses were more shape- than color-selective; neuronal responses were also relatively more shape-selective during discrimination than during fixation (paired t-test, p = 0.04). (B). Selectivity bias for neurons whose responses were well-fit by the color-dependent model. Relative selectivity of < 1 indicates most responses were more color- than shape-selective; the selectivity of each neuron was similar during fixation and discrimination (paired t-test, p = 0.07). (C). Selectivity bias for neurons whose responses were well-fit by the firing rate-dependent models. Responses of these neurons spanned the range from relatively more color-selective to relatively more shape-selective, and each neuron’s selectivity was similar during fixation and discrimination (paired t-test, p = 0.15).

In this representation, the upper right quadrant contains neurons that are relatively more shape- than color-selective in both tasks. In our population, most neurons in this quadrant are those whose responses are best explained by the shape-dependent model (**Fig. 7A**). Many of these neurons also fall significantly above the identity line (paired t-test using log-transformed ratios, p = 0.04), suggesting an increase in relative shape-selectivity during discrimination. In contrast, the lower left quadrant contains neurons that are relatively more color- than shape-selective in both tasks. In our population, most neurons in this quadrant are those whose responses are best explained by the color- dependent model (**Fig. 7B**). Although many of these neurons became more color- selective during behavioral engagement (points falling below the identity line), this trend was not significant (paired t-test using log-transformed ratios, p = 0.07).

Neurons whose responses are best explained by the linear and non-linear firing-rate dependent models span across the two quadrants (**Fig. 7C**), reflecting that responses of neurons in this group were not consistently more selective for either shape or color (see **Fig. 6**). Moreover, there was no significant change in selectivity bias in the two tasks (paired t-test using log-transformed ratios, p = 0.15).

#### Feature tuning in the population

Finally, we asked whether the feature tuning changed during discrimination trials for neurons best explained by feature- or firing rate-dependent models. To assess tuning changes, we constructed tuning curves and compared their width for fixation and discrimination trials. For example, we first computed shape-specific PSTHs by taking an average response for the 3 stimuli with the same shape, for a total of 3 shape-specific PSTHs. We then normalized and averaged the PSTHs across all neurons (see Methods for more details). Finally, we obtained 1 point from each PSTH by calculating the average response in the 50-200 ms window (as in Figs. 2B-4B), for a total of 3 points representing tuning. Since stimuli were tailored to each neuron, the tuning axis was arranged based on ranked order preference for shape, and responses were normalized relative to the response for the preferred shape. Color tuning curves were constructed equivalently, using color-specific responses. We averaged feature curves for neurons whose responses were best explained by each of the feature-dependent models and by either of the firing-rate dependent models, and compared tuning for discrimination and fixation trials. For neurons whose responses were best explained by firing rate or stimulus color, average feature tuning was similar. However, this was not the case for neurons whose response modulation was shape-dependent. While shape tuning during discrimination (red, **Fig. 8A**, left panel) was similar to that during fixation (black trace), color tuning (**Fig. 8A**, right) was relatively flatter during discrimination trials. These results are consistent with the findings from the feature selectivity analyses above, which suggested that shape-dependent neurons become relatively more shape-selective. Observations from the tuning curve analysis indicate that this significant change in feature selectivity may be the result of a loss of color selectivity, reflected in the broadening of color tuning in these neurons.

**Figure 8.**
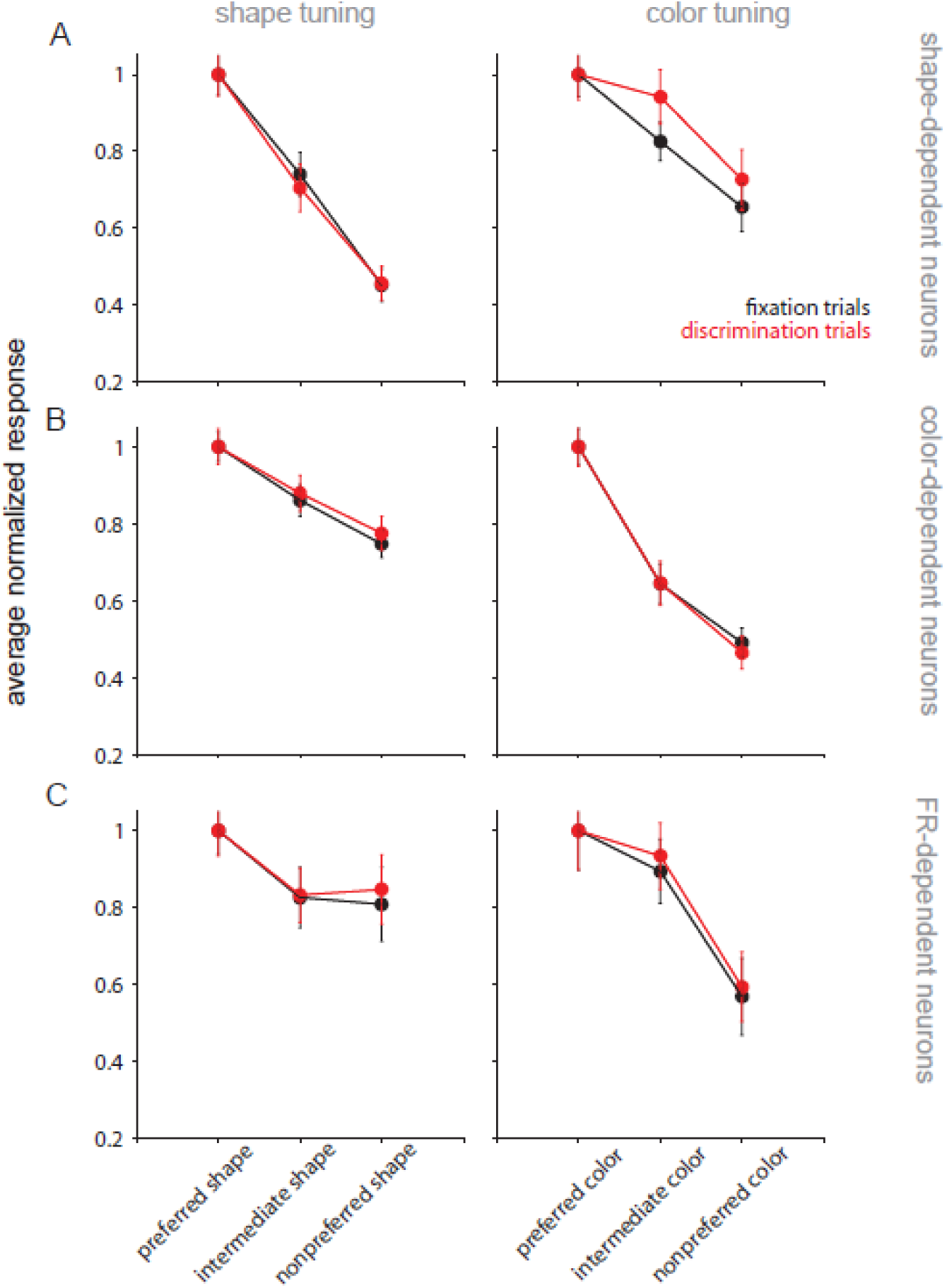
Feature tuning in the population. Shape (left column) and color tuning curves (right column) were constructed by taking average responses (as in Fig. 2B) from shape- and color-specific PSTHs for each neuron. In this figure we include the shape and color tuning of neurons whose responses are best described by the shape-dependent model (A), color-dependent model (B), and firing-rate dependent models (C). Feature-specific PSTHs were computed by taking, *e.g.*, average responses for different shapes across all colors. Before extracting average responses for tuning curves, PSTHs were normalized for each task, and averaged across groups of neurons. Bars show s.e.m.

In 9 neurons from Animal 1, we were able to obtain additional data from a discrimination task where animals judged stimulus color instead of stimulus shape. For 5/9 neurons, responses were best explained by the shape-dependent model, and for 4/9 by the color-dependent model. The responses of all 9 neurons were similar both in magnitude and selectivity for the two discrimination tasks. This observation suggests that the observed modulations are not related to which feature is judged in the discrimination task.

#### Influence of other behavioral factors

The results presented above suggest that for a subset of V4 neurons, a stimulus-dependent change can account for the difference in responses between fixation and discrimination trials. We examined whether two additional factors unique to the discrimination task contributed to the observed response modulation: the relationship between neuronal preference and the reference stimulus, and the saccadic eye movement reporting the animal’s decision.

#### Influence of reference stimulus

Previous studies of feature attention have reported that response modulation may be dictated by whether a neuron’s feature selectivity/preference for object features matches the stimulus feature being attended (Treue and Martinez-Trujillo, 1999; also see Bichot *et al*., 2005). For example, if a neuron’s preferred shape is a square, and the reference stimulus in the discrimination task is also square, that neuron’s responses would be enhanced; conversely, if a neuron’s non-preferred shape is a circle, its responses would be suppressed when the reference stimulus is a square. Although the present study compares two task conditions that are different than those typically used to study attentional effects, this interaction between stimulus and neuronal preference could contribute to the observed modulation in our data.

To address this question, we compared the responses in discrimination trials where the reference stimulus matched (reference-match) or did not match (reference-mismatch) the preferred shape and color of each neuron, relative to the overall response in the discrimination trials (see Eqn. 4a). This relative difference was very small for most neurons (median = 0.09). For a neuron with a mean firing rate of 100 spk/s, this value represents a mean stimulus response difference of <5 spk/s. The direction of this difference would be expected to be consistently positive, *i.e*. the reference-match trials should always be enhanced. The direction of response difference between reference-match and reference-mismatch trials was near 0 (median = −0.02; obtained by removing the absolute value operation from Eqn. 4a) and thus neurons in our population did not show a consistent enhancement of responses in reference-match trials.

Next, we examined whether tuning was similar in reference-match and reference-mismatch trials. We fit the linear and feature-dependent models separately for these subsets of data, and compared the regression coefficients (excluding the constant regression terms, as these additive terms would not reflect changes in tuning). The linear-gain model produced very similar fits for reference-match and reference-mismatch trials in all neurons (paired t-test of regression coefficients from ref-match and ref-mismatch data subsets, p = 0.6; correlation between coefficients, r = 0.99, p < 0.001). The coefficients from the shape- and color-dependent models were also similar between reference-match and reference-mismatch trials (paired t-test, p = 0.41 and p = 0.77, respectively). Overall, this result suggests that the relationship between reference stimulus identity and neuronal preference did not strongly contribute to the modulations we report above.

#### Influence of eye movement

Additionally, we considered whether the animal’s motor plan contributed to the observed modulation. In correct trials where the reference stimulus and the test stimulus had the same shape, the animal always performed a saccade to the target location on the right side of the screen; when the two stimuli had different shapes, the saccadic movement was to the target location on the left. Additionally, recording sites in both animals covered receptive field locations in the lower portion of the right visual hemifield; this imbalance could be reflected in the difference between right- and left-ward saccade trials. To determine whether the eye movement requirements of the task contributed to the effects we observed, we performed analyses similar to the reference match-mismatch above, except for data separated into two groups based on the saccade outcome of each discrimination trial.

First, we examined the magnitude of the neuronal responses in rightward-saccade and leftward-saccade trials (see Eqn. 4b). This response difference was very small compared to overall response rate for most neurons (median = 0.05), equivalent to a <3 spk/s response difference for a neuron with a mean firing rate of 50 spk/s. Next, we examined the direction of the difference between trials ending in rightward vs. leftward saccades (by removing the absolute value operation from Eqn. 4b). This value was distributed around 0 in the population (median = −0.02) suggesting that there was no consistent relationship between responses in trials ending in rightward vs leftward saccades. Thus, although some individual neurons showed some difference in response related to saccade direction, across the whole population there was no difference in response magnitude. Finally, we assessed differences in tuning by performing the linear and feature-based gain regressions separately on these data subsets, and comparing the coefficients for rightward- and leftward-saccade trials. These values were highly consistent for both models, indicating no change in tuning: for the linear-gain model, the Pearson correlation coefficient between regression coefficients for rightward and leftward trials was r = 0.999 (p < 0.001); for the shape-dependent gain model, r = 0.89 (p < 0.001); for the color-dependent gain model, r = 0.95 (p < 0.001). These results suggest that eye movements and the corresponding motor plan did not significantly contribute to the observed response modulations.

## DISCUSSION

We compared responses of 83 V4 neurons to colored shape stimuli in two task contexts: a passive fixation task and an active shape discrimination task. We found that most neurons displayed a change in responses between the two tasks. For 62/83 neurons in our population the modulation in responses during the discrimination task was best described on the basis of stimulus shape or color. Additionally, 3/83 neurons were well-fit by a nonlinear FR-based gain model; only 18/83 neurons were best fit by a linear FR-based gain model. Responses of the 20 neurons that were best fit by the shape-dependent models displayed a significant relative increase in shape selectivity during the discrimination task. Discrimination trial responses of these neurons were also associated with broadened color tuning.

### Relating observed trends to attention literature

Our observed changes in feature selectivity contrast with the broadly reported gain scaling of responses in sensory areas by attention (McAdams and Maunsell, 1999; Maunsell and Treue, 2006). Typically, attentional studies control stimulus presentation and cue the animal to attend to a relevant stimulus aspect (e.g. spatial location or orientation). Importantly, animal performs the same task in all attentional conditions. In contrast, the aim of our study was to compare visual encoding while an animal performs two different tasks. The behavioral task used here does not preclude effects of feature-based attention, and the goal of discriminating shape while ignoring color is similar especially to visual search studies (*e.g.* Mirabella *et al*., 2007). Top-down attentional mechanisms elaborated by previous literature are consistent with the observed response modulation in neurons that were firing-rate dependent (best fit by linear- and polynomial-gain models). The feature-dependent changes we observed in other neurons can arise via mechanisms which restructure neuronal preferences when the animal is engaged in a task, and which affect subpopulations of neurons with stronger feature-selectivity. The specificity of such a mechanism may be established in the network linking V4 to higher-level cognitive areas such as prefrontal cortex (Hamker, 2005; Gregoriou *et al*., 2009; Liebe *et al*., 2012), and may be developed or refined throughout the animals’ training to perform the discrimination task.

### Advantages of feature-dependent changes in response

When responses are feature-dependent during active behavior, selectivity bias changes: *e.g.*, responses of neurons best described by the shape-dependent model become relatively more shape-selective than color-selective during behavior. Since stronger feature selectivity is associated with higher discriminability along the feature dimension, the demands of the behavioral task may elicit response changes that increase the discriminability of the dimension best captured by a neuron’s responses (*e.g.* shape, for shape-selective neurons). This refinement of selectivity may be accomplished by selectively strengthening feature inputs, such as by thresholding, to enhance the strongest, or preferred, inputs to a cell, or vice versa - to suppress the less preferred inputs. In our analysis, the increase in relative shape selectivity in shape-dependent neurons appeared to be correlated with broadened color tuning; thus, the latter possibility may underlie the observed response changes.

It is difficult to conclude whether the tuning refinement observed in shape-dependent neurons improves shape coding without information about the noise covariance (Zhang and Sejnowski, 1999; Pouget *et al*., 1999); further studies involving simultaneous recordings from neuronal populations would help resolve this uncertainty. In a few sessions where the animal performed both shape discrimination and color discrimination tasks, the modulation in neuronal responses relative to the fixation task was the same regardless of the type of discrimination. This observation suggests that the observed response changes may be due to the gross differences between fixation and discrimination tasks, and not due to the behavioral relevance of the object feature. However, the animal from which these additional data were obtained had been trained extensively to judge object shape, and only later trained to judge object color. The animal’s training history may have contributed to two different behavioral goals eliciting the same response modulation.

### Relationship to other task-related factors

An existing model of feature attention links non-uniform scaling effects to the relationship between neuronal selectivity and the attended (or “reference”) stimulus in the behavioral task (the “feature-similarity model”; Martinez-Trujillo and Treue, 1999, 2004; Bichot *et al*., 2005). This model predicts enhanced responses when the attended stimulus matched the preferred stimulus of the neuron, and suppressed responses when the attended stimulus did not match. This effect leads to an increased signal-to-noise ratio and non-multiplicative scaling of the population response - that is, a narrowing in the width of the population tuning curve. In our data, we found that response modulation was not related to the feature similarity between the reference stimulus and the preference of the neuron.

In the discrimination task, the animals could respond at will after the test stimulus was presented (a reaction-time task), at which time the stimulus was immediately extinguished; this was unlike the fixation task, where the animal maintained its gaze for a fixed duration of stimulus presentation. Although reaction-time paradigms can introduce a confounding effect of eye movement planning, this did not contribute to the observed difference between the responses in the two tasks. Our analyses examined responses within a time window (50-200 ms from stimulus onset) well before the typical reaction time of the animals (260 ms). An additional influence of eye movement planning has been reported when an animal’s eye movement targets a location inside the receptive field, in which case V4 responses increase (conversely, targeting a location outside the receptive field suppresses responses; Han *et al*., 2009). Our recording sites targeted RFs only in the lower portion of the visual hemifield, which could have differentially affected responses on trials when the animal made a saccade to the target on the same side as the RF versus the target in the opposite hemifield. Moreover, “same” and “different” shape judgments in the discrimination task were always linked to a rightward- and leftward-directed saccade, respectively. Nevertheless, we found no relationship between eye movement and the response change during the discrimination task. This result may arise from two key differences: first, both of our targets lay outside the receptive field, so response changes consistent with previous findings would tend to be in the same direction and thus the magnitude of the difference would be smaller. Furthermore, the response changes reported by Han *et al*. were typically within a 50 ms epoch before the saccadic eye movement, and thus would largely be absent in the portion of the response we chose to analyze.

In summary, we have shown that responses of many V4 neurons are modulated by task context, and these changes in responses depend on the features of the stimulus.

## Acknowledgments

We thank Amber Fyall for expert animal care. This work was funded by NEI Grant R01EY018839 to A.P.; NEI Center Core Grant for Vision Research P30EY01730 to the University of Washington; NIH/ORIP Grant P51OD010425 to the Washington National Primate Research Center; and UW Vision Training Grant (NEI T32EY007031), NSF GRFP DGE-1256082, and UW Computational Neuroscience Training Grant (NIH T90DA032436) to D.V.P.

## Supplementary Figures and Text

**Figure S1.**
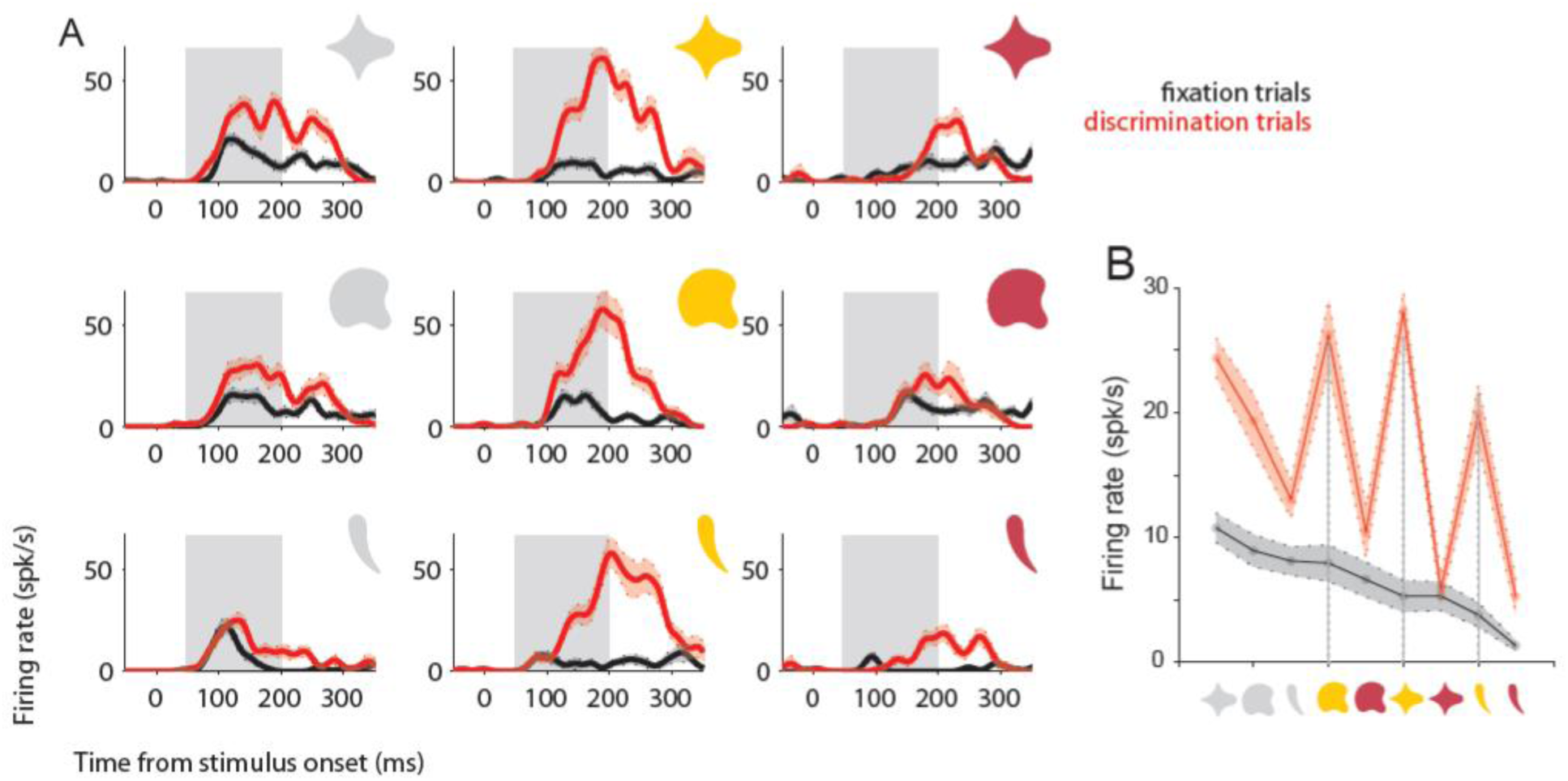
Responses of example neuron #4 (p150604) to stimuli in fixation and discrimination task conditions. Stimuli shown in light gray appeared white against the uniform gray background used in our experiment. (A). PSTH for responses of neuron #4 while the animal performed a fixation task or discrimination task; layout and color scheme as in Fig. 2A. Responses of neuron #4 were both color- and shape-selective during the fixation task (compare responses to stimuli in top row; stimuli in left column, respectively). Note that during the discrimination task, responses to a less preferred color (middle column) became stronger than the originally preferred color, and more similar in magnitude across different shapes. (B). Schematic representation of tuning for neuron #4, computed as in Fig. 2B. Grey dashed lines indicate responses to stimuli in middle column of (A).

Figure S1 shows responses of a neuron that appears to change stimulus preference in a manner consistent with a tuning peak shift. This neuron’s responses during fixation trials (black traces, Fig. S1A) were modulated by both shape and color (cf. PSTHs for stimuli in top row and left column, respectively). The stimulus shown in the top left panel elicited the strongest response from this neuron during fixation trials. During discrimination trials (red traces), firing rate increased for all stimuli. However, this neuron responded best to the stimulus in the top center panel during discrimination. Stimuli shown in the yellow color (center column) elicited a similar magnitude of responses during discrimination, although they evoked different responses during the fixation task.

This dependence is reflected in the S-R curve for this neuron (Fig S1B). During discrimination trials, stimuli with a yellow color elicited a larger increase in FR than expected from the ranked stimulus preference during fixation trials (*e.g.* dashed lines highlight responses to stimuli in center column of Fig. S1A). As for Neurons 2 and 3, responses of this neuron during discrimination trials were not consistent with a simple scaling of responses during fixation trials.

**Figure S2.**
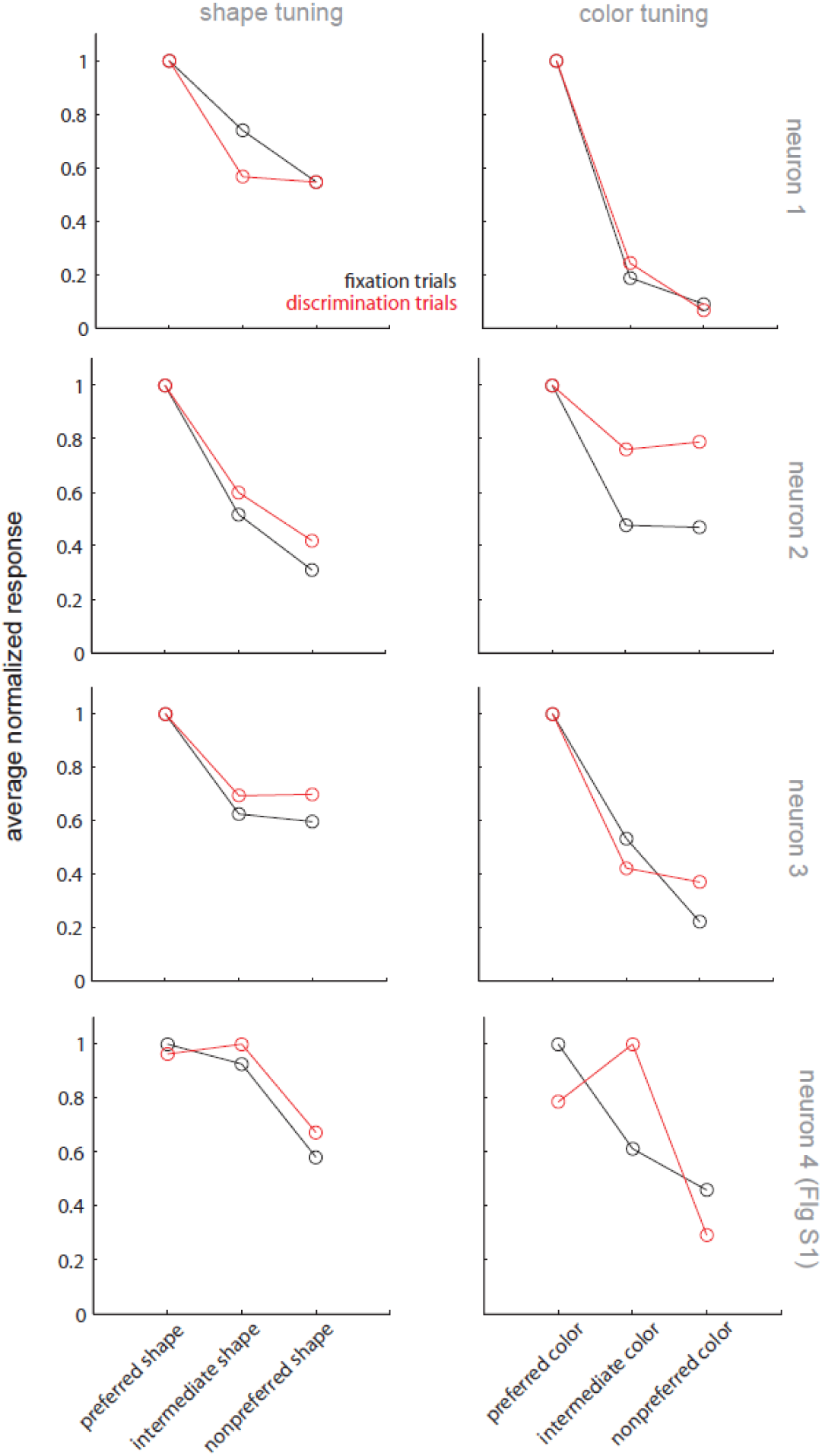
Feature tuning curves for example neurons 1-4. Shape (left column) and color tuning curves (right column) were constructed by taking average responses (as in Fig. 2B) from shape- and color-specific PSTHs for each example neuron, and then normalized by the response to the most preferred shape or color.

Figure S2 highlights that for Neurons 1-3, the most preferred stimulus shape or color remained the most preferred in both passive and active task contexts. This observation was representative of most of the neurons in our recorded population. Note also that color tuning for Neuron 2 broadens during discrimination, relative to fixation trials.

However, responses of Neuron 4 (bottom row; see also Fig. S1) show a change in the most preferred stimulus, especially for color. Such a change could be observed in just a few neurons.

